# Dissociable forms of uncertainty-driven representational change across the human brain

**DOI:** 10.1101/364638

**Authors:** Matthew R. Nassar, Joseph T. McGuire, Harrison Ritz, Joseph Kable

## Abstract

Environmental change can lead decision makers to shift rapidly among different behavioral regimes. These behavioral shifts can be accompanied by rapid changes in the firing pattern of neural networks. However, it is unknown what the populations of neurons that participate in such “network reset” phenomena are representing. Here we examined 1) whether and where rapid changes in multivariate activity patterns are observable with fMRI during periods of rapid behavioral change, and 2) what types of representations give rise to these phenomena. We did so by examining fluctuations in multi-voxel patterns of BOLD activity from human subjects making sequential inferences about the state of a partially observable and discontinuously changing variable. We found that, within the context of this sequential inference task, the multivariate patterns of activity in a number of cortical regions contain representations that change more rapidly during periods of uncertainty following a change in behavioral context. In motor cortex, this phenomenon was indicative of discontinuous change in behavioral outputs, whereas in visual regions the same basic phenomenon was evoked by tracking of salient environmental changes. In most other cortical regions, including dorsolateral prefrontal and anterior cingulate cortex, the phenomenon was most consistent with directly encoding the degree of uncertainty. However, in a few other regions, including orbitofrontal cortex, the phenomenon was best explained by representations of a shifting context that evolve more rapidly during periods of rapid learning. These representations may provide a dynamic substrate for learning that facilitates rapid disengagement from learned responses during periods of change.

## Introduction

Neural populations in rodent prefrontal cortex can undergo abrupt changes in firing concomitant with changes in performance in rule-based tasks (Durstewitz et al., 2010; Powell and Redish, 2016). Similar phenomena have been observed in the multi-voxel patterns in human fMRI data preceding changes in task strategy, leading to the notion that such changes might correspond to an “aha moment” at which the brain reorganizes to produce a new task set (Schuck et al., 2015). In rodent learning tasks that involve discontinuously changing reward contingencies, abrupt changes in firing of neurons in medial frontal cortex are observed more frequently during periods of uncertainty, during which animals appear to be searching for the best behavioral policy (Karlsson et al., 2012). It is unclear to what extent such phenomena are specific to medial frontal populations, or to what extent they might have an analog in human learning. Furthermore, while these “network resets” during periods of uncertainty are thought to play a role in behavioral flexibility in changing environments (Tervo et al., 2014) the exact computational role of abrupt changes in such neural representations remains unknown.

A number of different computational factors could explain previously observed network reset phenomena. First, and most simply, such abrupt changes would be expected in a neural representation of the current behavioral policy, which in some cases may be directly related to the motor program. Successful execution of learning requires maintenance and updating of a behavioral policy, which would tend to change more rapidly during periods of uncertainty.

Alternatively, reset phenomena might result from representation of higher-order computational variables used to appropriately calibrate the rate of learning. Recent work has highlighted a number of computational variables that are important for successful learning in the presence of discontinuous environmental changes (change points). In particular, humans tend to increase rates of learning according to the probability with which a given outcome reflects a change point in the behavioral contingency (*change*-*point probability*) and according to the relative imprecision of their estimate of the current contingency (*relative uncertainty*) (Nassar et al., 2010; 2012). These computational variables both increase following change-points, albeit with different dynamics, to mediate rapid incorporation of new information during and after periods of environmental change. Change-point probability and relative uncertainty correlate with BOLD responses across a wide swath of brain regions including some that jointly reflect both variables and some that uniquely reflect either change-point probability or uncertainty (McGuire et al., 2014). In principle, neural representations of either computational factor might involve patterns of activation that mimic “network reset” phenomena, yet this possibility has never been tested directly.

Another signal that might give rise to reset-like dynamics is a continuously evolving latent state representation. Latent states, which represent the relevant behavioral context in cases where it is not directly observable, can improve learning in the face of abstract stimulus categories or repeated episodes by efficiently partitioning learning across distinct behaviorally relevant contexts (Gershman and Niv, 2010). While previous work has focused primarily on the advantage of such representations for rapid reinstatement of previously learned behaviors (Gershman et al., 2010; Wilson et al., 2014), another advantage of such representations is that they could facilitate rapid disengagement from established behaviors that are no longer relevant. By appropriately partitioning data collected over time in a changing environment, such a mechanism could aid learning even if previously encountered environmental states to not recur. To accomplish this, such a latent state representation would need to evolve faster after a period of environmental change in order to effectively disengage from the previous behavioral context (Prescott Adams and MacKay, 2007; Wilson et al., 2010). While previous work has suggested that orbitofrontal cortex (OFC) might represent latent task states (Wilson et al., 2014; Schuck et al., 2016), it is unclear whether such representations transition dynamically during periods of rapid learning as would be necessary to efficiently mediate disengagement of learned responses that are rendered irrelevant by environmental change.

Here we examined whether and where uncertainty-linked network resets are observable in human fMRI data, and evaluated the most likely computational explanation for these phenomena in individual brain regions. We did so using a multistep approach. First, we identified signals that change rapidly from trial to trial during periods of uncertainty and rapid learning and potentially correspond to network resets (Karlsson et al., 2012). Second, we generalized this notion of representational change across pairs of non-consecutive trials using representational similarity analysis (RSA) (Nili et al., 2014). Third, we formalized a set of candidate computational explanations for network-reset phenomena and allowed these explanations to compete to explain multivariate brain activity (Kragel et al., 2018).

We observed rapid changes in multivariate activity patterns across widespread cortical regions during periods of uncertainty and rapid learning. Using RSA, we showed that patterns in motor regions were best described as reflecting behavioral policy, patterns of activation in occipital regions were best described as registering the occurrence of change-points, and patterns across much of the rest of the cortex appeared to reflect uncertainty. However, patterns of activation in a small number of regions including OFC were most consistent with dynamic latent state representations, suggesting a possible role for the OFC in translating learning signals into state changes that effectively disengage from behaviors learned in contexts that are no longer relevant.

## Results

To examine how neural signals change during periods of uncertainty we re-analyzed data from a previously published study that included recordings of fMRI BOLD signal and behavioral responses of human participants in a predictive inference task (McGuire et al., 2014). Participants played a video game in which they tried to get as many coins as possible (redeemable for money) by catching bags of coins dropped from a hidden helicopter in the sky. Thus, on each task trial, participants estimated the state of an unobservable variable (the position of a helicopter) based on the history of an observable variable (the position of bags dropped from that helicopter) (McGuire et al., 2014). The task included abrupt change points at which the position of the helicopter was resampled from a uniform distribution, which forced participants to rapidly revise beliefs about the helicopter location in order to maintain successful task performance. Here we refer to periods of consistent helicopter position as contexts (Fig 1a), such that the task could be described as requiring dynamic belief updating both within (Fig 1a; vertical) and across (Fig 1a; horizontal) contexts.

**Figure 1:**
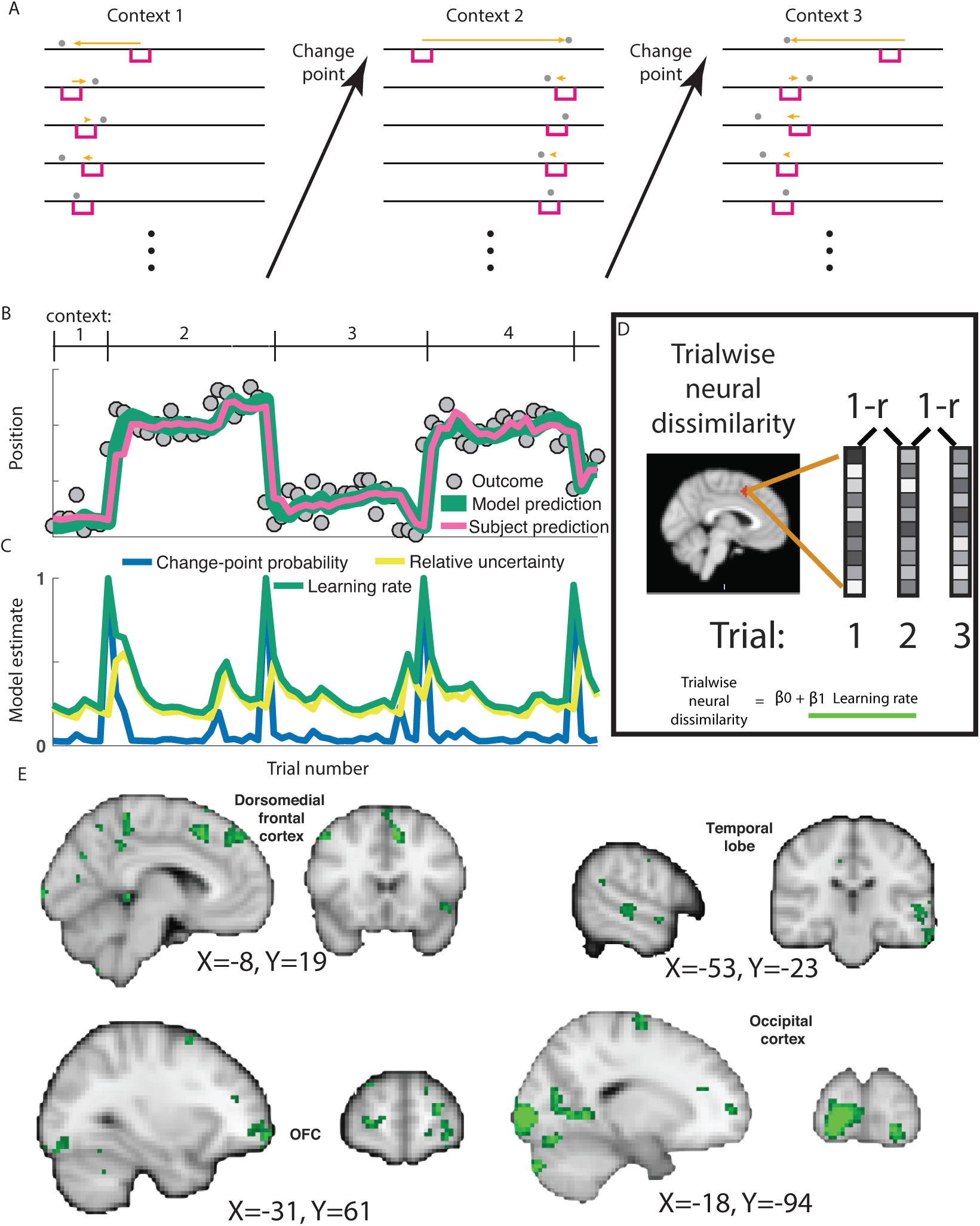
Trialwise neural dissimilarity is increased after change-points during periods of rapid learning for multiple brain regions. **A)** Participants were asked to move a bucket (pink rectangle) on each trial to the location most likely to deliver a reward (in the form of a bag containing coins). On each trial (stacked vertically) the participant would observe an outcome (bag location; gray circle) that they could use to update their bucket placement for the subsequent trial (orange arrow). Most contiguous trials were generated from the same context, which was defined by a fixed outcome distribution, however at occasional change points, the context (mean outcome location) shifted abruptly and unpredictably. **B)** An example sequence of outcomes (gray circles) and corresponding participant bucket placements (pink line) is plotted across trials. Participant bucket placements were well described by a normative learning model (green line) that adjusts learning rate according to change-point probability and relative uncertainty, which **(C)** are updated according to the model on each trial and evolve over time. **D)** Trial-wise measures of neural dissimilarity were computed on each trial as one minus the correlation coefficient between contiguous trial activations within a searchlight and regressed onto learning rates from the normative learning model to identify brain regions with BOLD activations that evolved more rapidly during periods of rapid learning. **E)** A diverse array of brain regions including occipital regions, dorsomedial prefrontal cortex, orbitofrontal cortex, and temporal regions displayed neural changes that were positively related to learning (green clusters). All images are thresholded at p = 0.001 uncorrected.

As we described in our previous report, adjustments in the rate at which participants revised beliefs in response to new information were well described by a normative learning model that adjusted learning according to two computational variables: change-point probability and relative uncertainty (Fig 1b, compare pink and green lines; (McGuire et al., 2014; Nassar et al., 2016)). Change-point probability reflects the Bayesian posterior probability that the helicopter has relocated on the current trial, and is largest on trials with large spatial prediction errors (Fig 1c, blue line). Relative uncertainty captures the degree to which uncertainty about the true helicopter location should drive learning, is greatest on the trial after a spike in change-point probability, and decays as a function of trials thereafter (Fig 1c, yellow line). Both of these factors affect the sensitivity of ongoing beliefs to new information (e.g., bag locations), which can be expressed in terms of a dynamic learning rate (Fig 1c, green). We sought to identify relationships between the sensitivity of behavior to incoming information (i.e., learning rate) and the sensitivity of neural representations to the same information.

The trial-to-trial dissimilarity in multivariate voxel activation patterns was related to the dynamic learning rates prescribed by the normative model (Fig 1d). Trial wise neural dissimilarity was computed for each pair of sequentially adjacent trials using a whole brain searchlight procedure and regressed onto an explanatory matrix that included model-based estimates of dynamic learning rates. A constellation of regions showed patterns of activation that changed more rapidly during periods of rapid learning after change points (Fig 1e). These regions included OFC, but also clusters in dorsomedial frontal cortex (DMFC), occipital cortex, and the temporal lobe. Thus, with a simple measure of representational change, we identified neural signals whose representations updated more rapidly during periods of learning in multiple brain regions (cf. (Karlsson et al., 2012)).

We next exploited representational similarity analysis (RSA) to extend and generalize the analysis above by incorporating information about the pairwise dissimilarity for all pairs of trials, not merely adjacent trial pairs. We hypothesized that the dissimilarity in neural representation for any pair of trials would depend on the cumulative amount of learning expected to occur between them under the normative model (see Methods). The hypothesized pattern of dissimilarity across trials is equivalent to what we would expect from a latent state representation that shifted rapidly at abrupt context transitions and concomitant periods of rapid learning, but remained relatively stable in periods when the statistics of the environment were stationary (Fig 2a). The pattern of dissimilarities predicted across adjacent trials using this strategy is exactly equivalent to the learning rates that served as the explanatory variable in the previous analysis (Fig 2b), but this generalization also makes predictions about the pattern of dissimilarities that would be observed across non-adjacent trials (Fig 2c). We used a searchlight to identify brain regions in which the neural dissimilarity matrix was positively associated with this hypothetical “shifting state representation” hypothesis matrix while controlling for fixed autocorrelation in the similarity structure (see Methods). A significant association was observed in a set of regions that overlapped with the results from the trial-wise dissimilarity analysis, including clusters in OFC, DMFC, occipital, and temporal regions (Fig 2d). As might be expected by the increased power owing to the non-adjacent trial comparisons afforded by RSA analysis, we also identified additional regions that were not clearly indicated by our previous analysis including a number of visual regions, left motor cortex, and bilateral hippocampus (Fig 2d).

**Figure 2:**
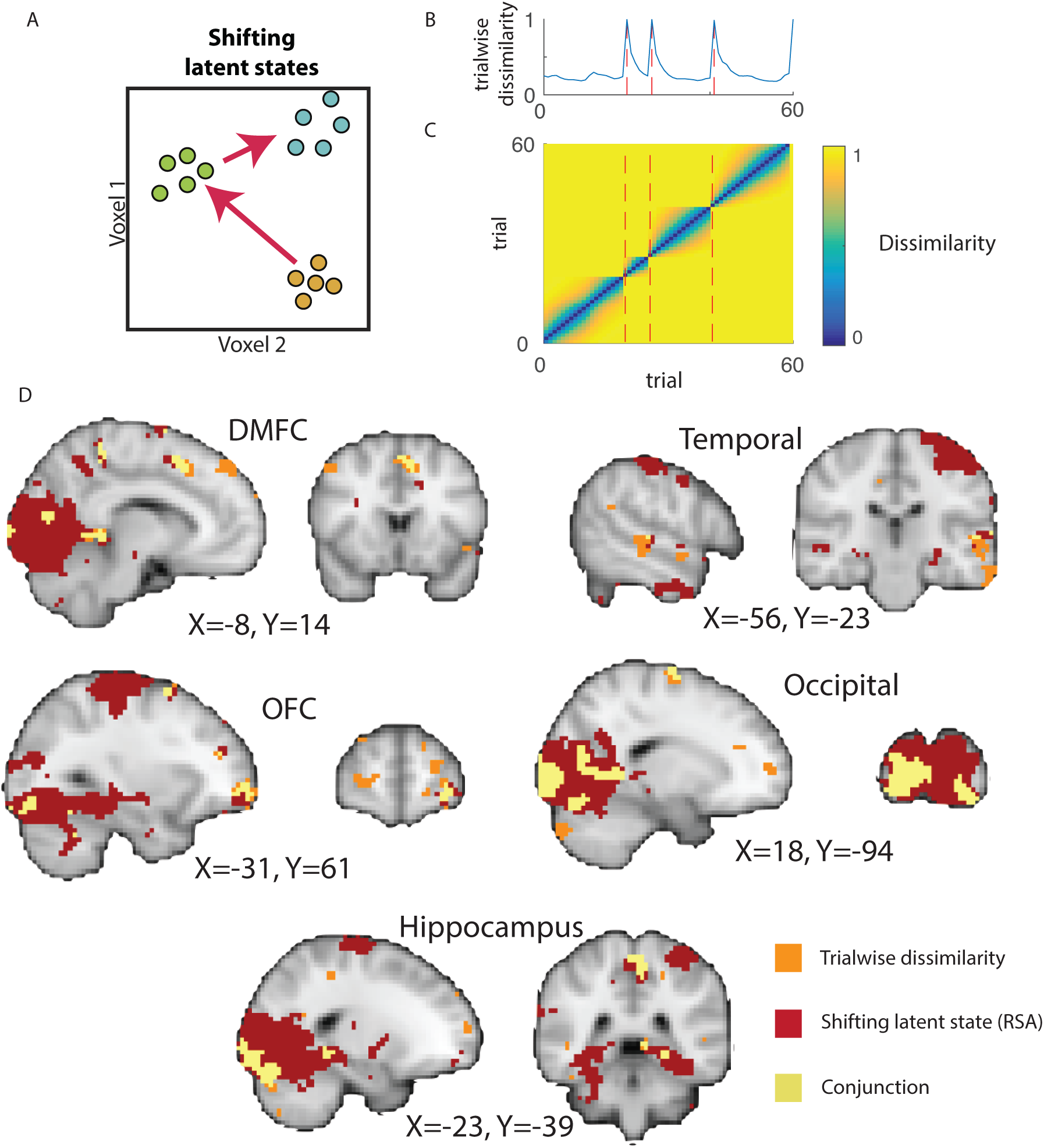
Representational similarity analysis reveals additional brain regions with representations that evolve more rapidly during periods of learning. **A)** In principle, rapid changes in neural representation coincident with learning might reflect a dynamic state representation that transitions rapidly at changes in context (see Fig 1a) and evolves more slowly as subjects develop accurate representations of the context. **B)** This would lead to greater trialwise dissimilarity immediately after change points in task context (blue line indicates simulated trialwise dissimilarity, red dashed lines indicate change points), but also to **(C)** unique patterns of dissimilarity across non-adjacent trials. **D)** A searchlight representational similarity analysis to identify such patterns revealed a constellation of regions (red) that overlapped substantially with that identified in the trialwise similarity analysis (orange; conjunction depicted in yellow), and also included additional regions such as left motor cortex, visual cortex, and hippocampus. All images are thresholded at p = 0.001 uncorrected.

We next sought to arbitrate among multiple possible causes for the varying rates of representational change. The rapid evolution of neural representations after change points might reflect different underlying computations in different brain regions. Our analysis focused on four candidate computations that could all theoretically drive network reset-like phenomena.

First, we considered the possibility that a brain region might reflect the behavioral policy of the participant. In our experimental task, the behavioral policy was reported directly by positioning a bucket at the predicted location (using a joystick) on each trial. For a given helicopter position, participants tended to place the bucket in a similar location, but changes in helicopter location corresponded to large changes in the bucket placement, which would correspond to abrupt transitions in a representation of behavioral policy after change points (Fig 3a). Occasionally, a new helicopter position was similar to one that had previously been encountered, such that a similar behavioral policy might be employed in two temporally separated contexts (Fig 3a; contexts 1&3).

**Figure 3:**
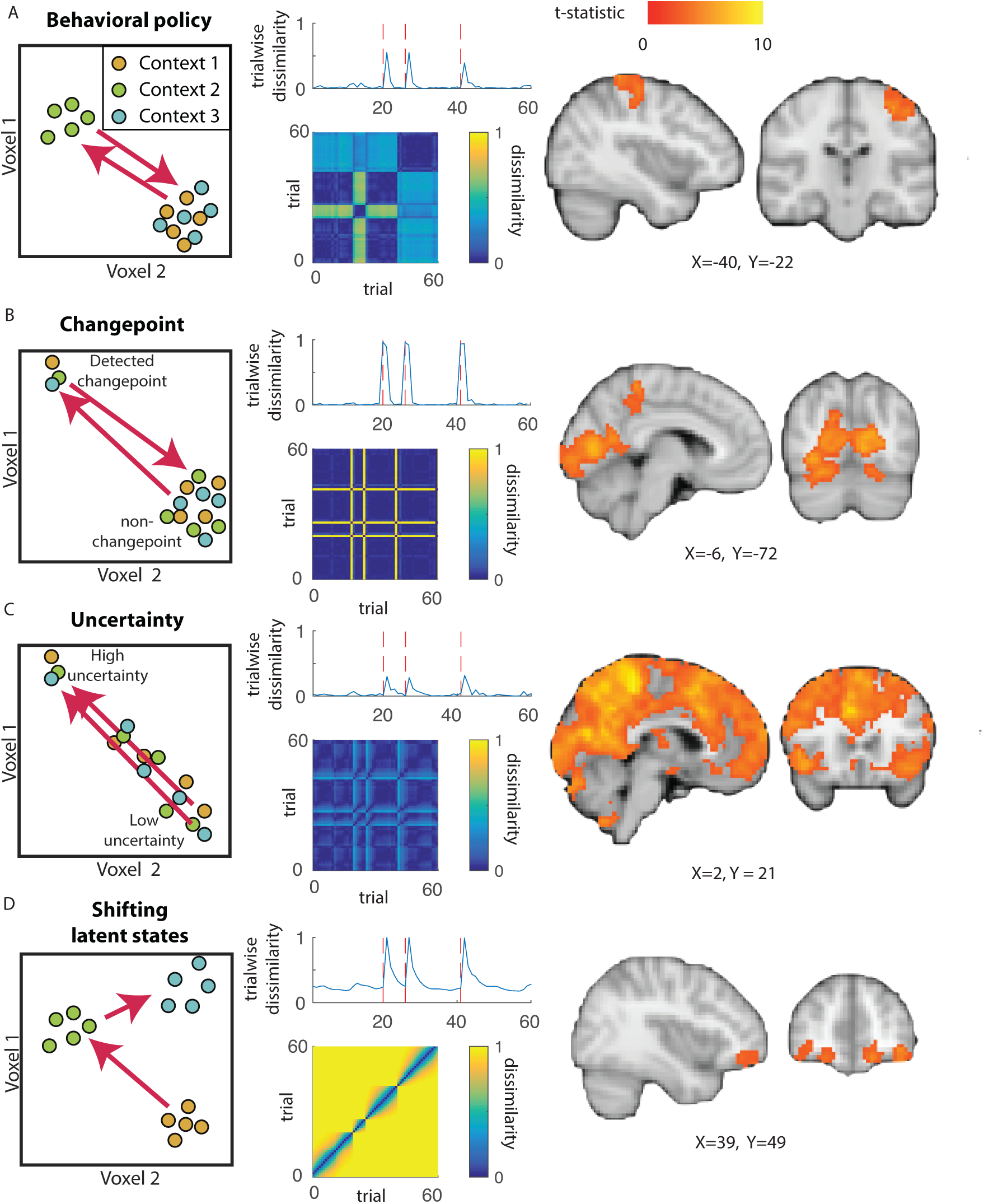
Dissociable explanations for task-driven changes in trialwise dissimilarity. *Left:* Context changes could affect different sorts of representations that are thought to be involved in task performance. A change in context could elicit a large representational change (arrows) in the behavioral policy (**A**), an internal assessment of change-point probability (**B**), the current level of relative uncertainty (**C**), or a latent state that shifts in proportion to learning (**D**). *Middle:* Each of these representations would predict increased trialwise dissimilarity after change points (top, red dotted lines indicate change points). However, dissimilarity matrices constructed across all trials (adjacent and non-adjacent) reveal unique representational profiles for each source of change-point related dissimilarity (bottom). *Right:* Patterns of voxel activations across trials revealed an anatomical dissociation between representations of behavioral policy (**A**; left motor cortex), change-point probability (**B**; occipital cortex), relative uncertainty (**C**; widespread), and shifting latent states (**D**; orbitofrontal cortex).

A second possible explanation for rapid representational change after change points is that the representations could reflect the current level of change-point probability or relative uncertainty. Change-point probability changes most dramatically at a change in the context (Fig 1c), leading to predicted trialwise neural dissimilarity time courses that do the same (Fig 3b). The level of relative uncertainty changes most rapidly immediately after change-points (Figure 1c), and a neural representation of relative uncertainty should do the same (Fig 3c). However, either of these representations should return to a fixed pattern for all epochs across the experimental session that share the same level of change-point probability or relative uncertainty, irrespective of the current helicopter position (Fig 3b–c).

A final computational explanation for rapid representational changes after change points is that such a signal may reflect a latent state that is used to partition learning across distinct contexts (Wilson et al., 2014). For example, each new helicopter position could be reasonably thought of as a new temporal context, during which learning from prior contexts should be discounted to minimize interference (Fig 1a). Since the helicopter position cannot be resolved exactly, such a context representation would be expected to evolve over time in proportion to the rate of learning about the current context. As described in Figure 2, this would lead to latent state representations that change rapidly at change points and immediately afterwards and change only minimally during periods of prolonged stability (Fig 3d). Unlike the other computational factors discussed above, a latent state representation would not necessarily exhibit any systematic similarity relation between one context and another – as our task did not include situations in which the helicopter returned exactly to a previously occupied position. Such a latent state signal might provide an evolving substrate to which outcomes could be linked in order to achieve rational adjustments of learning.

Each of these representations would yield more rapid changes in neural patterns after change points in our task, and indeed, they make very similar predictions for how neural dissimilarity metrics between adjacent trials should evolve over time (Fig 3 middle column, top plots). Predictions of trial-to-trial dissimilarity made for the four candidate computations were highly correlated (all average pairwise Pearson correlations [r] were greater than 0.45, with predictions for shifting latent representations particularly highly correlated with those for relative uncertainty [r = 0.80] and behavioral policy [r = 0.74]), suggesting that the representations of these computations could not be distinguished based on adjacent-trial dissimilarity alone.

However, the four candidate representations differed drastically in their predictions about the dissimilarity for non-adjacent pairs of trials. We constructed hypothesis matrices for each candidate representation by considering the expected difference in the computation of interest across all possible pairs of trials. These hypothesis matrices highlight qualitative features of each candidate computation; behavioral policy frequently undergoes abrupt shifts but often takes on a similar value to a previous state, change-point probability highlights differences between change point and non-change point trials, relative uncertainty highlights the differences between high relative uncertainty and other trials, and shifting latent states capture differences largely near the diagonal (Fig 3, middle column, bottom). Consistent with these qualitative differences, correlations between the hypothesis matrices for the different candidate representations were relatively low (all pairwise r < 0.16), suggesting that the candidate representations could be efficiently distinguished when considering the entire pairwise dissimilarity matrix.

We exploited these distinct predictions using a representational similarity analysis approach that allowed alternative explanations of representational change to compete to explain the observed neural dissimilarity matrix. Neural dissimilarity was computed for each pair of trials as one minus the spatial correlation of trial-activations across voxels in a searchlight and regressed onto an explanatory matrix that included the hypothesis matrices for all four candidate representations, along with a number of other explanatory terms designed to account for factors changing throughout the task and simple sources of variability such as autocorrelation (see Methods).

Representational similarity analysis supported distinct explanations for representational change in different anatomical regions. Behavioral policy provided a good description of BOLD activity patterns in left motor cortex (contralateral to the hand used to move the joystick and execute the behavioral policy) and visual cortex (Figure 3a, right; Table 1). Representations of change-point probability were prominent in occipital cortex and precuneus (Figure 3b; Table 1). Representations of relative uncertainty were widespread across the brain and included DMFC, dorsolateral prefrontal cortex, bilateral parietal cortices, insula, as well as some occipital and temporal regions (Figure 3c, right). Patterns of activation consistent with a latent state that shifts according to assessment of the current context were prominent in OFC and temporal cortex (Fig 3d, right; Table 1).

**Table 1:**
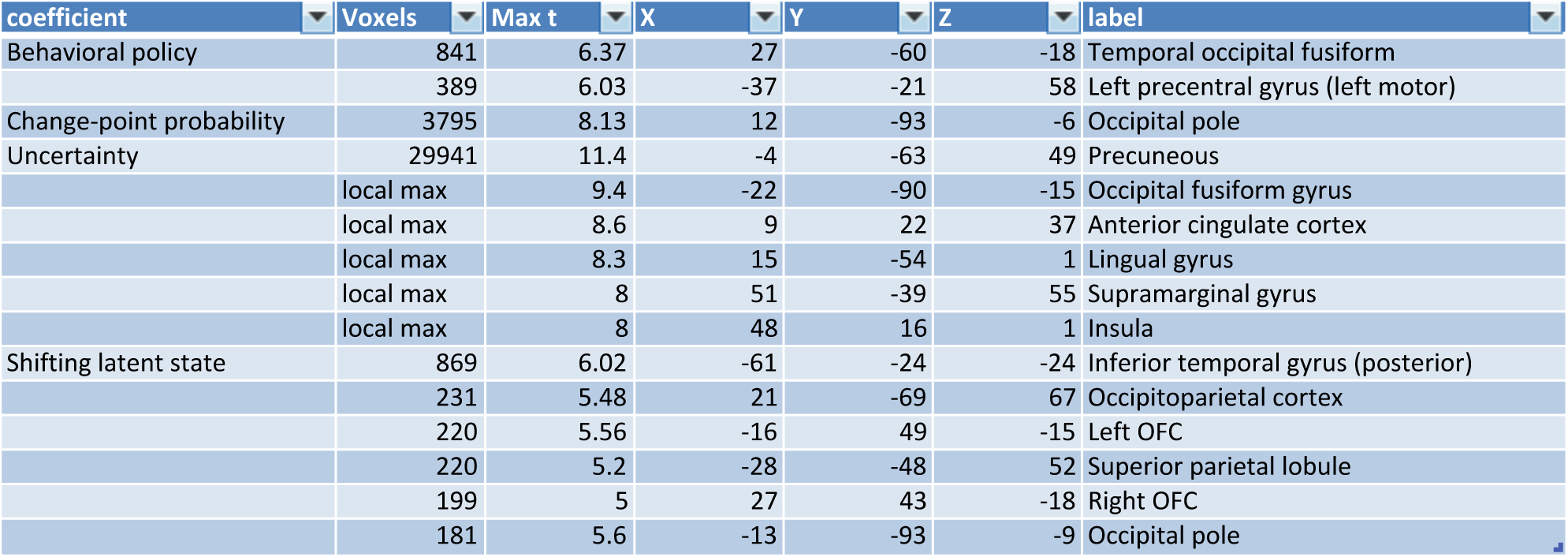
Peak voxel locations corresponding to behavioral policy, relative uncertainty, change-point probability and shifting latent state representations. Cluster size (in voxels), maximum (t-statistic) and MNI coordinates for each cluster surviving multiple comparisons correction.

The relationship between the neural dissimilarity in OFC and the dissimilarity structure predicted by a shifting latent state signal was robust to specific analysis choices. Patterns of activation in right and left OFC clusters were positively related to shifting latent state predictions in the context of our representational similarity regression analysis when using alternative pre-processing strategies such as omitting smoothing (Table 2) or including a spatial pre-whitening procedure (Table 3), both of which emphasize the high frequency components of the spatial pattern (Walther et al., 2016). The observed effects were not driven by relationships between additional explanatory variables included in the regression model, as exclusion of other explanatory variables yielded very similar relationships (Table 4). It is noteworthy that this was not true of all clusters that survived whole-brain correction in our representational similarity regression analysis; clusters identified in left superior parietal lobule and right occipital cortex were not related to the shifting latent state predictions in isolation (Table 4). Furthermore, the relationship between shifting latent state predictions and OFC patterns of activation was also robust to our assumptions about the exact timing of learning; a time shifted version of the shifting latent state hypothesis matrix that assumed learning occurred immediately upon observing a trial outcome could also describe similarity patterns observed in right and left OFC (Table 5).

**Table 2:** Latent state analysis with unsmoothed voxels. Regions-of-interest that showed a significant effect of shifting latent state, reanalyzed with unsmoothed voxels.

| Region | Mean Beta | t-value | p-value<br>(uncorrected) |
| --- | --- | --- | --- |
| Left inferior temporal gyrus | 0.0663 | 4.42 | 1.11e-4 |
| Left superior parietal lobule | 0.0491 | 3.51 | .00138 |
| Right occipital cortex | 0.1104 | 4.80 | 3.76e-5 |
| Left orbitofrontal cortex | 0.0541 | 3.36 | .00210 |
| Right orbitofrontal cortex | 0.0649 | 4.08 | 2.89e-4 |
| Left occipital pole | 0.0442 | 2.97 | .00574 |

**Table 3:** Latent state analysis with unsmoothed, pre-whitened voxels. Regions-of-interest that showed a significant effect of shifting latent state, re-analyzed with unsmoothed voxels that were spatial pre-whitened (Walther et al., 2016).

| Region | Mean Beta | t-value | p-value<br>(uncorrected) |
| --- | --- | --- | --- |
| Left inferior temporal gyrus | 0.0375 | 3.68 | 8.78e-4 |
| Left superior parietal lobule | 0.0175 | 1.82 | .0792 |
| Right occipital cortex | 0.0624 | 3.16 | .00347 |
| Left orbitofrontal cortex | 0.0256 | 2.27 | .0304 |
| Right orbitofrontal cortex | 0.0271 | 2.18 | .0367 |
| Left occipital pole | 0.0243 | 2.68 | .0116 |

**Table 4:** Minimal latent state analysis with unsmoothed voxels. Latent state effect in ROIs sensitive to latent state, re-analyzed with unsmoothed voxels and a model that only contained an intercept, the latent state predictor, and 15 off-diagonal autocorrelation terms.

| Region | Mean Beta | t-value | p-value<br>(uncorrected) |
| --- | --- | --- | --- |
| Left inferior temporal gyrus | 0.0693 | 4.57 | 7.37e-5 |
| Left superior parietal lobule | 0.0116 | 0.547 | .588 |
| Right occipital cortex | 0.0372 | 1.13 | .265 |
| Left orbitofrontal cortex | 0.0517 | 3.43 | .00172 |
| Right orbitofrontal cortex | 0.0586 | 3.93 | 4.45e-4 |
| Left occipital pole | 0.0539 | 3.47 | .00153 |

**Table 5:** Time shifted latent state analysis. Shifting latent state effect in ROIs sensitive to shifting latent state, re-analyzed using a time-shifted “shifting latent state” regressor in which representations at the time of outcome on a given trial are modeled as reflecting the beliefs that will guide behavior on the subsequent trial. This is offset by one trial from our original analysis, which assumed that representations upon viewing an outcome would reflect the beliefs that were formed in anticipation of that outcome, rather than the updated ones that incorporated it.

| Region | Mean Beta | t-value | p-value<br>(uncorrected) |
| --- | --- | --- | --- |
| Left inferior temporal gyrus | 0.0729 | 4.19 | 2.14e-4 |
| Left superior parietal lobule | 0.0656 | 4.22 | 2.00e-4 |
| Right occipital cortex | 0.0859 | 4.09 | 2.81e-4 |
| Left orbitofrontal cortex | 0.0720 | 3.98 | 3.91e-4 |
| Right orbitofrontal cortex | 0.0640 | 4.06 | 3.11e-4 |
| Left occipital pole | 0.0426 | 3.08 | .00435 |

In summary, while we found a number of regions that showed rapidly changing representations during periods of uncertainty following a context change, these reset-like phenomena were due to dissociable computational explanations. While a few regions were implicated in representing behavioral policy or change-point probability, most of these regions reflected relative uncertainty, and a smaller subset of regions including OFC were consistent with representing a latent state that is adjusted according to changes in context.

## Discussion

Neural representations in rodent medial frontal cortex rapidly change during periods of uncertainty (Karlsson et al., 2012). Here we demonstrate, in the context of a dynamic learning task, that such rapid representational changes are present in the BOLD signal in widespread cortical and subcortical regions. Furthermore, we showed that these rapid representational changes are consistent with several different computational explanations, which could be teased apart by considering the similarity structure of non-adjacent trials through representational similarity analysis.

Our analyses revealed distinct explanations for rapid representational changes in different brain regions. Focal representations of behavioral policy and change-point probability were identified in motor and visual cortex respectively, while widespread representations of relative uncertainty were observed throughout the brain. In addition, a small number of brain areas including the OFC had patterns of activation consistent with a form of shifting latent state representation that could speed disengagement from well-learned responses in a changing context.

Perhaps most straightforwardly, our analysis revealed that left motor cortex contained representations consistent with behavioral policy. In our task, this policy was completely concordant with the physical movement necessary to implement the behavioral policy. Thus, we interpret these results as a consequence of our experimental design, which required subjects to provide an analog behavioral output of their behavioral policy with their right hand on each task trial. Thus, this result was likely driven, at least in part, by a univariate effect of movement magnitude in the contralateral motor cortex.

Two other computations that we identified using this approach, change-point probability and relative uncertainty, had been the focus of a previous paper using this same dataset (McGuire et al., 2014). In the case of change-point probability, both univariate and RSA analyses revealed occipital cortex and precuneus as the locus of neural representation (see Figure 2c and (McGuire et al., 2014)). However, relative uncertainty representations identified using RSA were considerably more widespread than those identified through univariate activations (see Figure 2c and (McGuire et al., 2014)). This broader set of areas included some regions that were activated in the univariate analysis (e.g., DMFC), some that were deactivated in the univariate analysis (e.g., ventromedial prefrontal cortex), and some that were not identified in univariate analyses at all (e.g., temporal cortex). The near-ubiquitous cortical representation of relative uncertainty revealed by RSA is somewhat analogous to the widespread representations of reward prediction errors that have been identified using multivariate fMRI analysis methods (Vickery et al., 2011). Interestingly, both reward prediction errors and relative uncertainty have been suggested to be signaled through brainstem neuromodulatory systems that could potentially have widespread effects throughout the brain (Schultz, 1997; Yu and Dayan, 2005; Doya, 2008; Nassar et al., 2012).

In addition to providing a more sensitive tool to identify well-specified computational variables, RSA also allowed us to look for patterns of activity that could not easily be detected in univariate analyses. In particular, it allowed us to look for neural representations of a dynamically shifting state representation, without making strong assumptions about what the signal would look like at any given moment. It has been proposed that state representations provided by the OFC might serve to hasten learning in environments that include a small number of repeated contexts (Gershman and Niv, 2010; Wilson et al., 2014; Schuck et al., 2016). Here we hypothesized that shifts in the same state representations might implement the rapid learning that should and does follow change-points in outcome contingencies (Prescott Adams and MacKay, 2007; Nassar et al., 2010; Wilson et al., 2010). Such an implementation could make use of existing computational elements to efficiently partition learned associations that pertain to distinct and unrelated contexts, effectively creating the product partitions necessary for optimal inference amid change-points (Prescott Adams and MacKay, 2007).

In line with this idea, we identified signals in orbitofrontal cortex consistent with a shifting state signal that changed more rapidly during periods of learning. A neural population that encoded such a signal would be well positioned to transform a direct representation of dynamic learning rate, such as have been identified in cortical regions (Behrens et al., 2007; Krugel et al., 2009; McGuire et al., 2014) and thought to be broadcast through noradrenergic neuromodulation (Yu and Dayan, 2005; Nassar et al., 2012; Browning et al., 2015), into a proportional change in associative strength. Using a learning signal to control the rate of contextual shift could enable a simple associative neural network to accomplish the type of adaptive learning that has previously been modeled as a delta-rule update with a varying learning rate. In such a case, increases in apparent learning would be implemented through changes in the substrate for learning, or the active latent state, rather than by adjusting associative strength per se.

Representations of latent state that transition dynamically from one context to the next are similar in spirit to the concept of event segmentation in episodic memory (Ezzyat and Davachi, 2010). Segmenting events is useful in that it can allow memories that are embedded within the same event but separated in time to share associations, while memories that may be closer in time but embedded in separate events are maintained separately, preventing interference (Reynolds et al., 2007). One mechanism through which segmentation could be achieved involves dynamic adjustment of the time-constant in slowly fluctuating temporal context signals to effectively “reset” context at event boundaries (Howard and Kahana, 2002; Howard et al., 2010; Manning et al., 2011). Our data suggest a link between this aspect of episodic encoding and the dynamic adjustments of learning that have been observed at context boundaries (Behrens et al., 2007; Nassar et al., 2010; McGuire et al., 2014). However, aspects of our findings also raise questions about the extent of this link. While our results could be interpreted as supporting roles for OFC and temporal lobe in segmenting contexts, we did not observe the same phenomenon in the hippocampus, which is thought to play a key role in event segmentation (Ezzyat and Davachi, 2014; Hsieh et al., 2014; Shapiro, 2014). Instead, we found that representations in hippocampus, like many other brain regions, were best explained as representing uncertainty itself. One potentially relevant detail is that previous contexts were not systematically re-visited in our task, reducing demands for episodic retrieval. An interesting avenue for future work would be to examine how the representations we identified respond when the context abruptly returns to a previously encountered state, such as might require a form of mental time travel for successful performance (Manning et al., 2011).

Our results, especially regarding the OFC, demonstrate the utility of analyzing the representational similarity of multi-voxel patterns of activity in concert with computational modeling. Such an approach allowed us to identify neural representations consistent with a specific computational role for OFC, which in principle could not have been isolated in our task with univariate activation or multivariate classification analyses.

In summary, we show that shifts in the statistics of the environment during a dynamic learning task induced both elevated learning and changes in neural representation. These changes in neural representation were attributed to specific computations using RSA. Our results identified widespread representations of relative uncertainty throughout the brain, together with more focal representations of change-point probability and behavioral policy. In addition, a small number of brain areas including the OFC had patterns of activation consistent with a shifting latent state representation that could speed unlearning of irrelevant information in a changing context.

## Methods

### Behavioral task and analysis

For details of the behavioral task and data analysis, see our previous report (McGuire et al., 2014). Briefly, 32 human subjects performed a computerized predictive inference task in an MRI scanner while undergoing functional neuroimaging. Each trial required the subject to move a bucket across the horizontal axis of a screen (starting from a “home position” at the right-hand edge, using a joystick controlled by the right hand) to a location that they believed most likely to be underneath a helicopter that was occluded by clouds and thus not directly observable. On each trial, the helicopter would drop a bag that contained either high value or neutral items. Bag locations were normally distributed and centered on the helicopter location (incentivizing bucket placement under the inferred helicopter location). On the majority of trials (90%) the helicopter would remain in the same location as in the previous trial, but occasionally (10%) the helicopter would relocate to a new position along the horizontal axis of the screen (selected randomly and uniformly).

### MRI data acquisition and preprocessing

T1-weighted MPRAGE structural images (0.9375 × 0.9375 × 1mm voxels, 192 × 256 matrix, 160 axial slices, TI=1100ms, TR=1630ms, TE=3.11ms, flip angle=15°), T2^∗^-weighted EPI functional data (3mm isotropic voxels, 64 × 64 matrix, 42 axial slices tilted 30° from the AC-PC plane, TR=2500ms, TE=25ms, flip angle=75°), and fieldmap images (TR=1000ms, TE=2.69 and 5.27ms, flip angle=60°) were acquired on a 3T Siemens Trio with a 32 channel head coil. Functional data were acquired in 4 runs, each of which lasted 9 minutes and 25 seconds (226 images).

Data were preprocessed using AFNI (Cox, 1996; 2012) and FSL (Jenkinson et al., 2002; Smith et al., 2004; Jenkinson et al., 2012) in the following steps: 1) slice timing correction (AFNI’s *3dTshift*), 2) motion correction (FSL’s *MCFLIRT*), 3) fieldmap-based geometric undistortion, alignment with structural images, and registration to the MNI template (FSL’s *FLIRT* and *FNIRT*), 4) spatial smoothing with a 6mm FWHM Gaussian kernel (FSL’s *fslmaths*), 5) outlier attenuation (AFNI’s *3dDespike*), and intensity-scaling by a single grand-mean value in each run (FSL’s *fslmaths*). The resulting functional time series was deconvolved to estimate trial activations at the time of the bag drop using the least squares-separate method (Mumford et al., 2012) implemented in Matlab.

### Multivariate fMRI analysis

Multivariate analyses were conducted in spherical searchlights (radius = 3 voxels) across the entire brain. Within each searchlight, the neural dissimilarity between each pair of trials was computed as one minus the spatial Pearson correlation between the voxel-wise activations for those trials.

Trial-to-trial dissimilarity scores were extracted by extracting the i=j-1 diagonal elements from the dissimilarity matrix, which corresponded to the dissimilarity between adjacent trials (see Figure 1d). The dissimilarity scores were regressed onto an explanatory matrix containing an intercept, and dynamic learning rates prescribed by a normative learning model, yielding one coefficient of interest per subject, per searchlight. Dynamic learning rates were estimated as the sum of change-point probability and relative uncertainty minus their product (see Figure 1c; (Nassar et al., 2016)). These latent variables were estimated with a parameter-free normative model that took subject prediction errors as an input according to the following set of recursive equations:

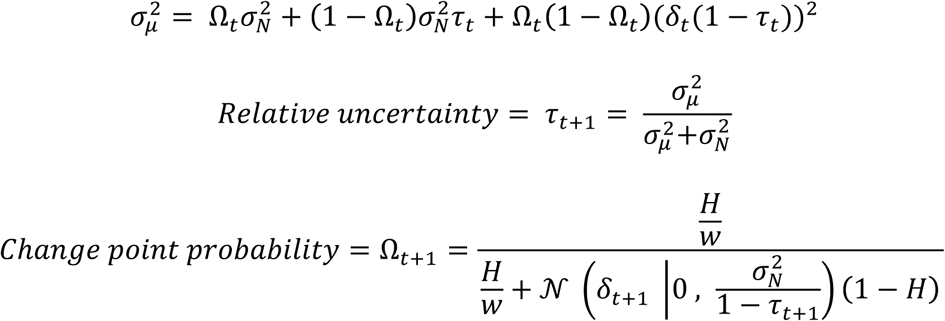

where 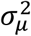 is the total variance in beliefs about the helicopter location (the generative mean), 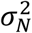 is the variance in the distribution of outcomes (bag drops) around that mean, *δ_t_* is the prediction error, and *H* is the hazard rate and *w* is the width of the screen. For a full derivation of the model and terms see (Nassar et al., 2010) and for a complete description of the method for estimating latent variables see (Nassar et al., 2016).

In general, change-point probability and relative uncertainty were both increased after change-points, albeit with different latencies, leading to learning rates that decay slowly as a function of time within context. Learning rates quantifying sensitivity to information provided on trial j were aligned with the trial-to-trial dissimilarity between trials j and j+1. Thus, our analysis targeted patterns of activity whose degree of change between trials j and j+1 reflected normative learning predicted to occur from the outcome presented on trial j. The first 3 trials from each block were removed from analysis as they occurred at the onset of fMRI acquisition.

Trial-to-trial dissimilarity analysis described above could be thought of as a special case of the general idea that the similarity between each pair of trials might be inversely related to the learning done between them. Because this pattern of similarity is what might be expected to emerge from a representation of the latent task state, which transitions abruptly from one context to the next and remains relatively stable after many trials in a well learned context, we will refer to it as the shifting latent state dissimilarity matrix. The hypothesis matrix for shifting latent states was generated by computing the extent to which the inference on trial i would factor into the inference on trial j, assuming normative learning:

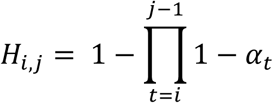

where H is the shifting latent state dissimilarity matrix and α is the learning rate prescribed by a normative model (Nassar et al., 2010), such that more prescribed learning between two trials corresponded to higher values of α, a smaller product term, and thus a greater dissimilarity. The i=j-1 diagonal of this matrix is 1-(1-α_t_), or just α_t_, and thus equivalent to the vector of trial-to-trial dissimilarities described above. However, the shifting latent state hypothesis matrix also includes information about other elements in the matrix, potentially offering a more powerful construct to ask a similar question. We examined whether this similarity structure was reflected in the neural dissimilarity between trials in each spherical searchlight. The lower triangle of the neural dissimilarity matrix was regressed onto a hypothesis matrix that included an intercept, the shifting latent state hypothesis matrix (lower triangle), and 15 dummy variables designed to remove the influence of autocorrelation on the coefficient of interest. These autocorrelation terms were derived from 15 off-diagonal binary matrices in which a single off diagonal (i = j-1; i = j-2; i = j-3… i = j-15) was set to one. These matrices were constructed to account for any variance in the neural dissimilarity matrices that could be explained by a fixed signal autocorrelation. To be sure that autocorrelation could not affect our analysis of interest, we also set all elements of the shifting latent state similarity matrix that fell outside of this range (trials separated by more than fifteen trials) to the maximum dissimilarity value.

To better understand the computations that give rise to rapid changes in neural patterns during periods of learning after a helicopter relocation, we constructed an exhaustive set of hypothesis matrices and conducted a representational similarity analysis in which these representations could compete to explain structure in neural dissimilarity matrices. This analysis required generating hypothesis matrices for various factors that could relate to task uncertainty, learning, or explain nuisance variance in the dissimilarity matrices. Hypothesis matrices were generated for three additional explanatory variables of interest: 1) subject prediction (behavioral policy), 2) relative uncertainty, 3) change-point probability. We also included six additional nuisance variables: 4) the bag drop’s location, 5) signed prediction error (ie, the distance between the prediction and the bag drop), 6) high CPP [to account for patterns of activity that may asymmetrically encode CPP], 7) high RU [to account for patterns of activity that may asymmetrically encode RU], 8) outcome reward value, and 9) task block. For factors 1-5 and 8, element (i,j) of the hypothesis matrix corresponded to the absolute difference in that factor on trials i and j. For factor 9, dissimilarity values were set to 0 for trials in the same block and 1 for trials in different blocks. Dissimilarity matrices for factors 6 & 7 were computed as one minus the multiplicative interaction of the model variable (6=change-point probability, 7=relative uncertainty) on trials i and j, such that similarity was only hypothesized when the model-derived term took on a high value on both trials. These terms allowed the model to capture asymmetric representations of the two factors governing learning in our model, such as a representation that converged for values of high relative uncertainty but did not show any consistent pattern of activation when relative uncertainty was low.

The lower triangle of the neural dissimilarity matrix was extracted and regressed onto an explanatory matrix consisting of an intercept and the lower triangle of all hypothesis/nuisance matrices (including the shifting latent state and nuisance autocorrelation terms), yielding one coefficient per variable, per subject, per searchlight (Chikazoe et al., 2014; Kragel et al., 2018). Group level analyses were conducted by computing t-statistics across subjects for each variable and searchlight. Cluster-based permutation testing using cluster mass with a cluster forming threshold of p<0.001 and an alpha of 0.01 was used to identify significant activations (Nichols and Holmes, 2002).

## Reference

Behrens TEJ, Woolrich MW, Walton ME, Rushworth MFS (2007) Learning the value of information in an uncertain world. Nature Neuroscience 10:1214–1221.

Browning M, Behrens TE, Jocham G, O’Reilly JX, Bishop SJ (2015) Anxious individuals have difficulty learning the causal statistics of aversive environments. Nature Neuroscience 18:590–596.

Chikazoe J, Lee DH, Kriegeskorte N, Anderson AK (2014) Population coding of affect across stimuli, modalities and individuals. Nature Neuroscience 17:1114–1122.

Cox RW (1996) AFNI: software for analysis and visualization of functional magnetic resonance neuroimages. Comput Biomed Res 29:162–173.

Cox RW (2012) AFNI: what a long strange trip it’s been. NeuroImage 62:743–747.

Doya K (2008) Modulators of decision making. Nature Neuroscience 11:410–416.

Durstewitz D, Vittoz NM, Floresco SB, Seamans JK (2010) Abrupt Transitions between Prefrontal Neural Ensemble States Accompany Behavioral Transitions during Rule Learning. Neuron 66:438–448.

Ezzyat Y, Davachi L (2010) What Constitutes an Episode in Episodic Memory? Psychol Sci 22:243–252.

Ezzyat Y, Davachi L (2014) Similarity Breeds Proximity: Pattern Similarity within and across Contexts Is Related to Later Mnemonic Judgments of Temporal Proximity. Neuron 81:1179–1189.

Gershman SJ, Blei DM, Niv Y (2010) Context, learning, and extinction. Psychological Review 117:197–209.

Gershman SJ, Niv Y (2010) Learning latent structure: carving nature at its joints. Current Opinion in Neurobiology 20:251–256.

Howard MW, Kahana MJ (2002) A Distributed Representation of Temporal Context. Journal of Mathematical Psychology 46:269–299.

Howard MW, Shankar KH, Jagadisan UKK (2010) Constructing Semantic Representations From a Gradually Changing Representation of Temporal Context. Top Cogn Sci 3:48–73.

Hsieh L-T, Gruber MJ, Jenkins LJ, Ranganath C (2014) Hippocampal Activity Patterns Carry Information about Objects in Temporal Context. Neuron 81:1165–1178.

Jenkinson M, Bannister P, Brady M, Smith S (2002) Improved optimization for the robust and accurate linear registration and motion correction of brain images. NeuroImage 17:825–841.

Jenkinson M, Beckmann CF, Behrens TEJ, Woolrich MW, Smith SM (2012) FSL. NeuroImage 62:782–790.

Karlsson MP, Tervo DGR, Karpova AY (2012) Network resets in medial prefrontal cortex mark the onset of behavioral uncertainty. Science 338:135–139.

Kragel PA, Kano M, Van Oudenhove L, Ly HG, Dupont P, Rubio A, Delon-Martin C, Bonaz BL, Manuck SB, Gianaros PJ, Ceko M, Reynolds Losin EA, Woo C-W, Nichols TE, Wager TD (2018) Generalizable representations of pain, cognitive control, and negative emotion in medial frontal cortex. Nature Publishing Group.

Krugel LK, Biele G, Mohr PNC, Li S-C, Heekeren HR (2009) Genetic variation in dopaminergic neuromodulation influences the ability to rapidly and flexibly adapt decisions. Proceedings of the National Academy of Sciences 106:17951–17956.

Manning JR, Polyn SM, Baltuch GH, Litt B, Kahana MJ (2011) Oscillatory patterns in temporal lobe reveal context reinstatement during memory search. Proceedings of the National Academy of Sciences 108:12893–12897.

McGuire JT, Nassar MR, Gold JI, Kable JW (2014) Functionally dissociable influences on learning rate in a dynamic environment. Neuron 84:870–881.

Mumford JA, Turner BO, Ashby FG, Poldrack RA (2012) Deconvolving BOLD activation in event-related designs for multivoxel pattern classification analyses. NeuroImage 59:2636–2643.

Nassar MR, Bruckner R, Gold JI, Li S-C, Heekeren HR, Eppinger B (2016) Age differences in learning emerge from an insufficient representation of uncertainty in older adults. Nature Communications 7:11609.

Nassar MR, Rumsey KM, Wilson RC, Parikh K, Heasly B, Gold JI (2012) Rational regulation of learning dynamics by pupil-linked arousal systems. Nature Neuroscience 15:1040–1046.

Nassar MR, Wilson RC, Heasly B, Gold JI (2010) An approximately Bayesian delta-rule model explains the dynamics of belief updating in a changing environment. Journal of Neuroscience 30:12366–12378.

Nichols TE, Holmes AP (2002) Nonparametric permutation tests for functional neuroimaging: a primer with examples. Hum Brain Mapp 15:1–25.

Nili H, Wingfield C, Walther A, Su L, Marslen-Wilson W, Kriegeskorte N (2014) A Toolbox for Representational Similarity Analysis Prlic A, ed. PLoS Comput Biol 10:e1003553.

Powell NJ, Redish AD (2016) Representational changes of latent strategies in rat medial prefrontal cortex precede changes in behaviour. Nature Communications 7:12830.

Prescott Adams R, MacKay DJC (2007) Bayesian Online Changepoint Detection. eprint arXiv:07103742:–.

Reynolds JR, Zacks JM, Braver TS (2007) A computational model of event segmentation from perceptual prediction. Cogn Sci 31:613–643.

Schuck NW, Cai MB, Wilson RC, Niv Y (2016) Human Orbitofrontal Cortex Represents a Cognitive Map of State Space. Neuron 91:1402–1412.

Schuck NW, Gaschler R, Wenke D, Heinzle J, Frensch PA, Haynes J-D, Reverberi C (2015) Medial Prefrontal Cortex Predicts Internally Driven Strategy Shifts. Neuron 86:331–340.

Schultz W (1997) A Neural Substrate of Prediction and Reward. Science 275:1593–1599.

Shapiro ML (2014) Time and Again. Neuron 81:964–966.

Smith SM, Jenkinson M, Woolrich MW, Beckmann CF, Behrens TEJ, Johansen-Berg H, Bannister PR, De Luca M, Drobnjak I, Flitney DE, Niazy RK, Saunders J, Vickers J, Zhang Y, De Stefano N, Brady JM, Matthews PM (2004) Advances in functional and structural MR image analysis and implementation as FSL. NeuroImage 23 Suppl 1:S208–S219.

Tervo DGR, Proskurin M, Manakov M, Kabra M, Vollmer A, Branson K, Karpova AY (2014) Behavioral Variabilitythrough Stochastic Choice and Its Gating by Anterior Cingulate Cortex. Cell 159:21–32.

Vickery TJ, Chun MM, Lee D (2011) Ubiquity and Specificity of Reinforcement Signals throughout the Human Brain. Neuron 72:166–177.

Walther A, Nili H, Ejaz N, Alink A, Kriegeskorte N, Diedrichsen J (2016) Reliability of dissimilarity measures for multi-voxel pattern analysis. NeuroImage 137:188–200.

Wilson RC, Nassar MR, Gold JI (2010) Bayesian online learning of the hazard rate in change-point problems. Neural Comput 22:2452–2476.

Wilson RC, Takahashi YK, Schoenbaum G, Niv Y (2014) Orbitofrontal cortex as a cognitive map of task space. Neuron 81:267–279.

Yu AJ, Dayan P (2005) Uncertainty, neuromodulation, and attention. Neuron 46:681–692.

